# Ataxin-2 regulates the synaptonemal complex to ensure chromosome pairing during female meiosis

**DOI:** 10.64898/2026.02.11.705321

**Authors:** Vernon L. Monteiro, Zhuyi Wang, Jose F. Santé, Camilla Roselli, Baskar Bakthavachalu, Mani Ramaswami, Craig Smibert, Thomas R. Hurd

## Abstract

Sexual reproduction relies on meiotic recombination and the accurate segregation of homologous chromosomes to generate viable, genetically diverse gametes. While the molecular mechanisms of recombination and chromosome segregation are well studied, the upstream regulatory cues that drive expression of key meiotic genes remain poorly understood, especially in metazoans. Emerging evidence suggests that post-transcriptional regulation plays a central role in initiating and coordinating the meiotic program in both fruit flies and mammals. Here, we identify the RNA-binding protein Ataxin-2 (Atx2) as a crucial regulator of meiosis in *Drosophila melanogaster*. We show that Atx2 positively regulates meiotic factors, especially components of the synaptonemal complex (SC), a structure essential for pairing, recombination and segregation of homologous chromosomes. In *Atx2*-depleted germ cells, SC component mRNA and protein levels are markedly reduced, leading to defective SC assembly and maintenance. Consequently, homologous chromosomes fail to pair properly, which is essential to ensure homolog segregation and prevent aneuploidy. These findings uncover Atx2 as a key regulator of the SC and highlight an underappreciated layer of gene regulation essential for accurate meiotic chromosome segregation and fertility.

## INTRODUCTION

Accurate segregation of homologous chromosomes at the first meiotic division is essential for Mendelian inheritance, and failures in this process are a major cause of miscarriage in humans (Llano and Pendás 2023; Adams and Davies 2025; Williams and Hawley 2025; Zickler and Kleckner). A key structure ensuring faithful homolog segregation is the synaptonemal complex (SC), a conserved structure that aligns homologous chromosomes along their length (Williams and Hawley 2025). In *Drosophila*, loss of SC function compromises homolog alignment leading to segregation errors during meiosis (Williams and Hawley 2025; Zickler and Kleckner). Accordingly, proper SC assembly is essential for accurate meiotic chromosome segregation and reproductive success.

Transcriptomic analyses have revealed that many meiotic genes are transcribed prior to meiosis and are post-transcriptionally regulated (Martin et al. 2022; Vallés et al. 2024). This includes genes encoding SC components, even though full SC assembly onto chromosomes occurs only at the onset of meiosis (Williams and Hawley 2025). Tight regulation of SC protein abundance and is essential, as SC components have a strong intrinsic tendency to self-assemble into aberrant structures that can disrupt gametogenesis if not properly controlled (Goldstein 1987; Hughes and Hawley 2020). While recent studies have identified post-translational mechanisms controlling SC dynamics, much less is known about how SC component expression and synthesis are regulated.

We recently identified the RNA-binding protein Ataxin-2 (Atx2) as a candidate post-transcriptional regulator acting at the mitotic-to-meiotic transition (Palozzi et al. 2022; Monteiro et al. 2025). We found that Atx2 is required for the inheritance of healthy mitochondria, a process temporally coupled to meiotic entry (Palozzi et al. 2022). However, defects in mitochondrial quality control alone would be expected to compromise offspring fitness rather than cause complete sterility. The requirement for Atx2 is fertility in both *D. melanogaster* and *C. elegans* (Satterfield et al. 2002; Ciosk et al. 2004; Maine et al. 2004) therefore suggests additional roles for Atx2 in female gametogenesis. Across species, Atx2 family members promotes the stability and translation of their target mRNAs by recruiting polyadenylation complexes and/or interacting with poly(A)-binding proteins (PABPs) (Mangus et al. 1998; Satterfield and Pallanck 2006; Yokoshi et al. 2014; Lee et al. 2017; Inagaki et al. 2020; Singh et al. 2021; Nadimpalli et al. 2022; Petrauskas et al. 2024). Because Atx2 acts at the developmental window of meiotic entry and functions as a positive regulator of mRNA stability and translation, we hypothesized that Atx2 directly promotes meiotic entry by post-transcriptionally upregulating meiotic mRNAs.

Here, we show that Atx2 indeed promotes the accumulation of key meiotic transcripts, including those encoding SC components. Loss of Atx2 results in reduced SC mRNA and protein levels, impaired SC assembly and maintenance, and defects in chromosome clustering and homolog pairing, processes essential for accurate homolog segregation and the prevention of aneuploidy. Together, these findings reveal Atx2 as a key regulator of the SC complex and meiotic progression and suggest a critical role for post-transcriptional control in ensuring reproductive fidelity in *Drosophila*.

## RESULTS

### Identification of Atx2 target genes

Across species, Atx2 has a well-established role in promoting the stability and translation of target mRNAs (Mangus et al. 1998; Satterfield and Pallanck 2006; Yokoshi et al. 2014; Lee et al. 2017; Inagaki et al. 2020; Singh et al. 2021; Nadimpalli et al. 2022). Thus, to gain insight into Atx2’s function during meiosis, we characterized the transcriptomes of control and *Atx2*-depleted cells using RNA sequencing. Because Atx2 also functions in other cell types in *Drosophila* (Satterfield and Pallanck 2006), we first specifically isolated meiotic germ cells to ensure that our analysis captured Atx2-dependent transcript regulation in this developmental context.

In the Drosophila ovary, female germ cells undergo four mitotic divisions in region 1 before entering meiosis in region 2 (Figure 1A and B). To isolate meiotic germ cells, we used a transgenic reporter expressing GFP under the control of the *bam* promoter (*bamP*) (Chen and McKearin 2003). Although *bamP*-GFP is expressed in mitotically dividing germ cells, the GFP persists into early meiotic stages, which are far more abundant than mitotic germ cells, making it a useful marker for enriching early meiotic germ cells. We performed fluorescence-activated cell sorting (FACS) on *bamP*-GFP ovaries (Figure 1A–D) in control and *Atx2* knockdown conditions. We confirmed the efficacy of *Atx2* knockdown by immunofluorescence on GFP-Atx2 (Figure 1E).

**Figure 1.**
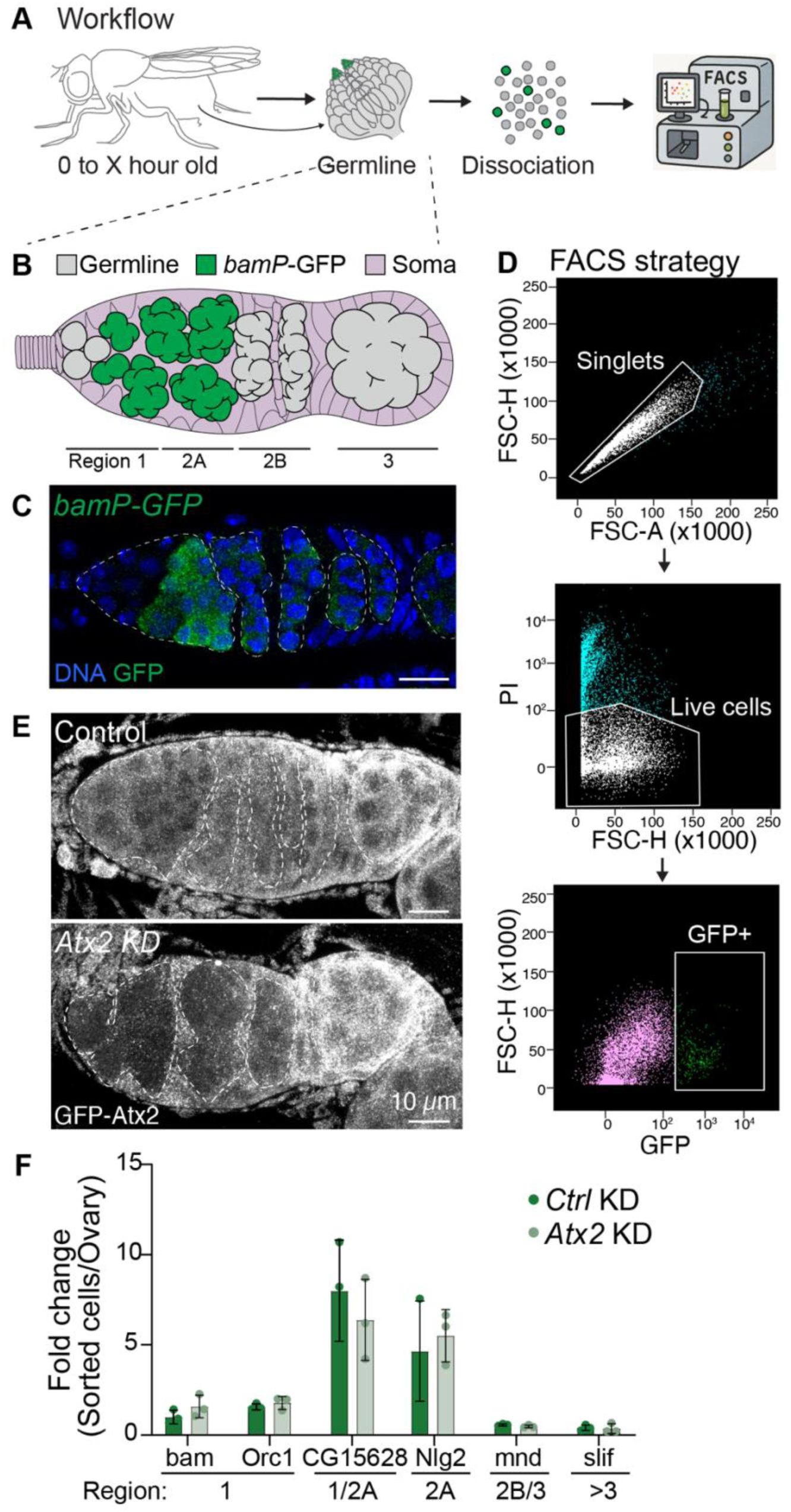
Isolation of meiotic germ cells from control and *Atx2*-depleted ovaries. **(A)** Schematic of workflow. **(B)** Schematic of female Drosophila germarium subdivided into regions 1, 2A and 2B, and 3 based on developmental stages. **(C)** Confocal micrograph of a germarium expressing GFP driven by the *bam* promotor and stained with DAPI (DNA, blue). Scale bar, 10 µm. White dashed line, germline. **(D)** FACS strategy to isolate *bamP*-GFP+ germ cells. **(E)** Confocal micrograph of a germarium expressing GFP-Atx2 expressing control (mCherry) shRNA or *Atx2* shRNA in the germline stained with anti-GFP. Scale bar, 10 µm. White dashed line, germline. **(F)** RT-qPCR analysis of stages specific markers in control and *Atx2*-depleted isolated germ cells relative to whole ovaries. Data are means ± s.d. of three independent replicates.

qPCR analysis of FACS-isolated *bamP*-GFP positive germ cells confirmed strong enrichment for marker genes expressed at the mitotic-to-meiotic transition (such as *CG15682*) and immediately afterward during the early stages of meiosis (such as *Nlg2*) (Slaidina et al. 2021) (Figure 1F). In contrast, markers expressed prior to meiosis (e.g., *bam*, *Orc1*) or at later stages of oogenesis (e.g., *mnd*, *slif*) were not strongly enriched (Slaidina et al. 2021), indicating that the isolated population consisted predominantly of early meiotic germ cells.

Using these isolated meiotic germ cells, we performed RNA sequencing on control and *Atx2*-depleted samples (Figure 2). We extracted total RNA and enriched for mRNA using ribosomal RNA depletion and primed cDNA synthesis using random primers. We performed RNA sequencing on at least three biological replicates per condition. Because chorion and vitelline membrane transcripts originate from somatic contamination and can dominate the dataset due to high expression (Spradling and Mahowald 1980), we removed them prior to gene normalization and downstream differential-expression analyses. Principal component analysis showed strong reproducibility across biological replicates and clear clustering by genotype (Supplementary Figure 1).

**Figure 2.**
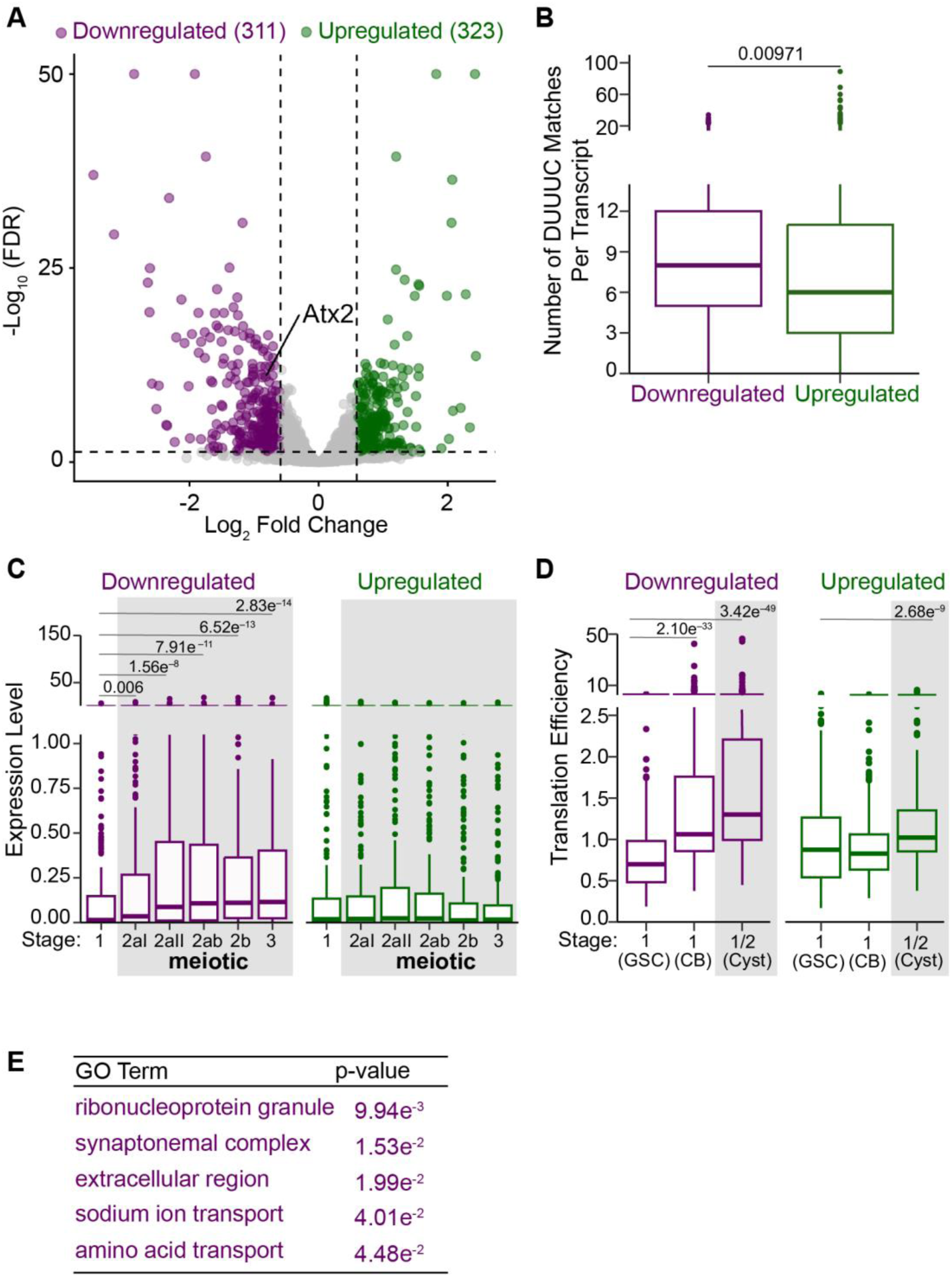
Atx2 increases the levels of transcripts involved in meiosis and sexual reproduction. **(A)** Volcano plot showing differential gene expression between FACS-isolated meiotic germ cells from control and Atx2-depleted ovaries. Significantly downregulated genes are shown in purple and significantly upregulated genes in green. Y-axis limited to -Log_10_(50) (see Table S1 for complete data). **(B)** Putative Atx2 targets are enriched for the Atx2-binding DUUUC motif (Santos et al. 2025). **(C)** Expression of putative Atx2 targets (genes downregulated upon Atx2 depletion) and of upregulated genes across distinct stages of germline development: prior to meiotic entry (stage 1), at the onset of meiotic entry (stages 2aI and 2aII), and during meiotic progression (stages 2ab, 2b, and 3). Expression data are from single-cell RNA sequencing (Slaidina et al. 2021). **(D)** Translation efficiencies of putative Atx2 targets (genes downregulated upon *Atx2* depletion) and of upregulated genes across distinct stages of germline development: prior to meiotic entry (stage 1 GSC, *nosGal4>UAS-tkv* and cystoblast (CB), *nosGal4>bam RNAi*), and subsequent differentiation into cysts (*nosGal4>bam RNAi, hs-bam*). Translation efficiencies are from polysome sequencing (Martin et al. 2022; McCarthy et al. 2022). **(F)** Gene Ontology (GO) term enrichment analysis for genes upregulated (B) and downregulated (C) upon Atx2 depletion. Five representative driver GO terms based on significance and non-redundancy are displayed.

We focused our analysis on transcripts decreased upon *Atx2* depletion because of Atx2’s established role in stabilizing mRNAs (Mangus et al. 1998; Satterfield and Pallanck 2006; Yokoshi et al. 2014; Lee et al. 2017; Inagaki et al. 2020; Singh et al. 2021; Nadimpalli et al. 2022). Differential expression analysis using DESeq2 identified 311 transcripts that were significantly downregulated in *Atx2*-depleted cells (|fold change| > 1.5; false discovery rate < 0.05) (Figure 2A; Supplementary Table 1). To determine whether these downregulated transcripts represent direct Atx2 targets, we tested for enrichment of the known Atx2 RNA-binding DUUUC motif (where D=A, G or U) (Yokoshi et al. 2014; Singh et al. 2021; Santos et al. 2025) in transcripts reduced upon *Atx2* depletion, relative to upregulated transcripts, which are unlikely to represent direct targets. The DUUUC motif was significantly enriched among downregulated transcripts (Figure 2B). These results indicate that transcripts reduced upon *Atx2* depletion are enriched for direct Atx2 targets and suggest that, in meiotic germ cells, Atx2 recognizes its targets through a DUUUC motif.

We next examined how putative Atx2 targets behave during normal germline development compared with non-targets. Using published RNA-sequencing and polysome-profiling datasets spanning key stages before, during, and after meiotic induction (Slaidina et al. 2021; Martin et al. 2022; McCarthy et al. 2022), we found that Atx2 targets show a marked increase in RNA abundance as germ cells transition from mitosis to meiosis, with levels remaining high thereafter (Figure 2C). In contrast, transcripts upregulated upon *Atx2* depletion showed little or no change across these developmental stages (Figure 2C). Consistent with the RNA trends, Atx2 targets also exhibited a larger increase in translation efficiency in more differentiated stages than upregulated genes (Figure 2D). Together, these findings reinforce a model in which Atx2 positively regulates the translation of mRNAs and possibly their stability at the onset of meiosis.

Lastly, to identify functional themes among Atx2-regulated genes, we performed Gene Ontology (GO) enrichment analysis. Putative Atx2 targets (i.e., those downregulated upon *Atx2* knockdown) were significantly enriched for GO terms associated with ribonucleoprotein granules, the SC, and the germ plasm (Figure 2E; Supplementary Table 2), consistent with Atx2 promoting meiotic entry. Among the ribonucleoprotein granule–associated genes were multiple RNA-binding proteins with essential roles in oogenesis and meiosis, including *nanos* (Slaidina and Lehmann 2014) and *orb* (Barr et al. 2019). Notably, we identified nearly all core components of the SC among Atx2 targets, which we focus on in subsequent analyses. Together, these results are consistent with a central role for Atx2 in promoting germline differentiation and meiotic progression through post-transcriptional regulation of key meiotic transcripts.

### Atx2 upregulates synaptonemal complex components

In *Drosophila*, several core components of the SC have been defined (Figure 3A), and our RNA sequencing data indicated that mRNA levels for all core SC genes, except for *cona* and the cohesion-associated factor *ord*, were reduced upon *Atx2* knockdown relative to control knockdowns (Figure 3B). To validate that Atx2 promotes SC transcript accumulation, we examined *c(3)G* mRNA, which encodes the essential transverse filament of the SC, *in vivo* using hybridization chain reaction fluorescent *in situ* hybridization (HCR-FISH). In heterozygous control ovaries, *c(3)G* mRNA was abundant in germline cysts prior to and during the onset of meiosis (Figure 3C, arrows). In contrast, *c(3)G* mRNA was markedly reduced in *Atx2* null trans-heterozygotes (Figure 3C and D), consistent with our RNA sequencing data and indicating that Atx2 is required to maintain *c(3)G* transcript levels in early meiosis. We next assessed C(3)G protein and found a pronounced reduction in *Atx2* mutant ovaries compared with controls (Figure 3E, arrows; and 3F), mirroring the loss of *c(3)G* mRNA and consistent with a potential role for Atx2 in stabilizing C(3)G mRNA and upregulating its translation. Together, these data are consistent with a role for Atx2 in upregulating SC component mRNAs and proteins at the onset of meiosis.

**Figure 3.**
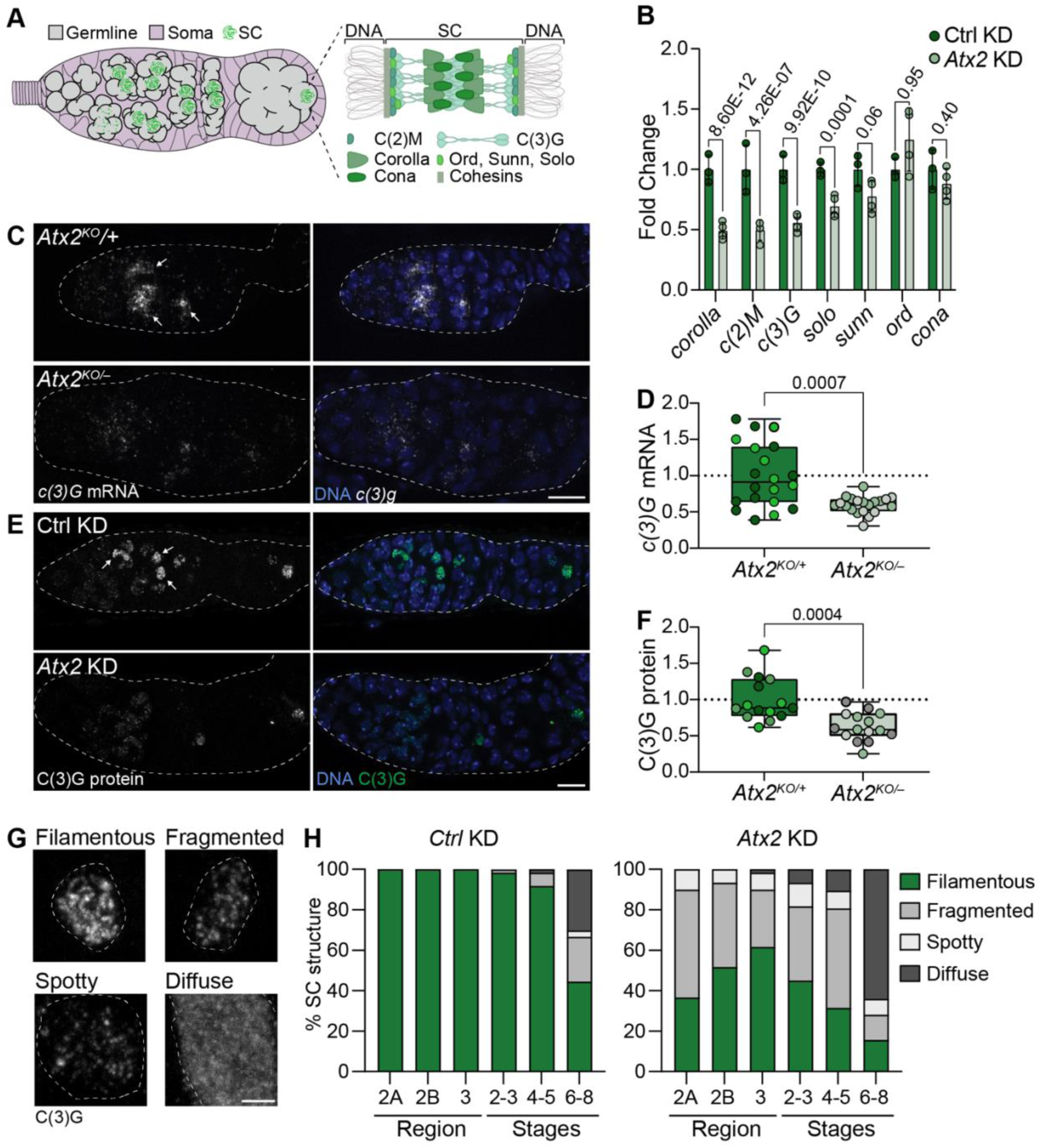
Atx2 promotes the SC. **(A)** Schematic of female Drosophila germarium showing the staging and timing of SC expression and assembly. **(B)** Normalized fold change RNA levels of SC components in control and *Atx2*-depleted germ cells from RNA sequencing experiments. p-values, Benjamini-Hochberg adjusted. **(C)** Confocal micrograph of germaria from heterozygous control and germline ablated *Atx2* ovaries. *c(3)G* mRNA was detected using HCR-FISH (white) and DNA stained with DAPI (blue). Scale bar, 10 µm. Dashed line, germarium. Arrows, *C(3)G* mRNA signal. **(D)** Quantification of (**C**). Fold change of *c(3)G* fluorescence. Data are means ± s.d. of two independent replicates (10 germaria per replicate). *p*-values, unpaired Welch’s t-test. **(E)** Confocal micrographs of germaria expressing control (mCherry) shRNA or *Atx2* shRNA in the germline stained with anti-C(3)G serum and DAPI. Scale bar, 10 µm. Dashed line, germline. Arrows, C(3)G signal. **(F)** Quantification of (**E**). Fold change of C(3)G fluorescence. Data are means ± s.d. of three independent replicates (3-6 germaria per replicate). *p-values*, unpaired Welch’s t-test. **(G)** Confocal micrographs of oocyte nuclei stained with anti-C(3)G serum representing the different SC phenotypes observed. SC structure in control (mCherry) germline knockdown (top left) and *Atx2* germline knockdowns (top right, bottom left and right). White dashed line, nuclei. Scale bar represents 2 µm. **(H)** Quantification of the SC throughout development in control and *Atx2* germline depleted ovaries.

### Atx2 is essential for synaptonemal complex formation and maintenance

Since core components of the SC are reduced in *Atx2-*depleted germ cells, we speculated that SC assembly and/or maintenance might be impaired. Although SC component genes are transcribed prior to meiosis, their loading along chromosome arms only becomes detectable after meiotic entry in region 2A (Hughes et al. 2018). At this stage, SC is typically observed in up to four nuclei within the 16-cell cyst; however, as the cyst development proceeds, three of these nuclei disassemble their SC and exit the meiotic program, leaving a single pro-oocyte in region 3 that retains fully assembled SC along its chromosomes (Hughes et al. 2018). SC only begins to disassemble in the pro-oocyte during mid-prophase (stage 5) and is fully absent by stages 7 to 9, marking the transition from mid to late prophase (Hughes et al. 2018).

To assess SC formation throughout oogenesis, we stained control and *Atx2*-depleted ovaries with the SC marker C(3)G, classifying SC morphology into four categories: filamentous (wild-type), fragmented, spotty, and diffuse (Figure 3G). In control ovaries, filamentous SC persisted until approximately stage 5, after which it progressively disassembled, displaying increasingly fragmented, spotty, and diffuse morphologies (Figure 3H). In contrast, *Atx2* knockdown ovaries showed reduced levels of the SC (see Figure 2) and impaired SC formation from the outset, with most nuclei exhibiting fragmented SC morphology (Figure 3H). This phenotype modestly improved by region 3, although 40% of cysts still showed abnormal SC, but worsened again from stage 2 onward. These observations indicate that Atx2 is critical for the formation and maintenance of a wild-type SC, likely through its role in upregulating the SC.

### Atx2 is required for proper centromere pairing and clustering

During meiosis, homologous chromosomes must first pair at their centromeres and then cluster with other paired homologs (Takeo et al. 2011; Tanneti et al. 2011; Christophorou et al. 2013; Joyce et al. 2013) (Figure 4A). In *Drosophila*, SC components are required for both centromere pairing and clustering. Given Atx2’s role in promoting SC assembly, we asked whether loss of Atx2 disrupts these processes.

**Figure 4.**
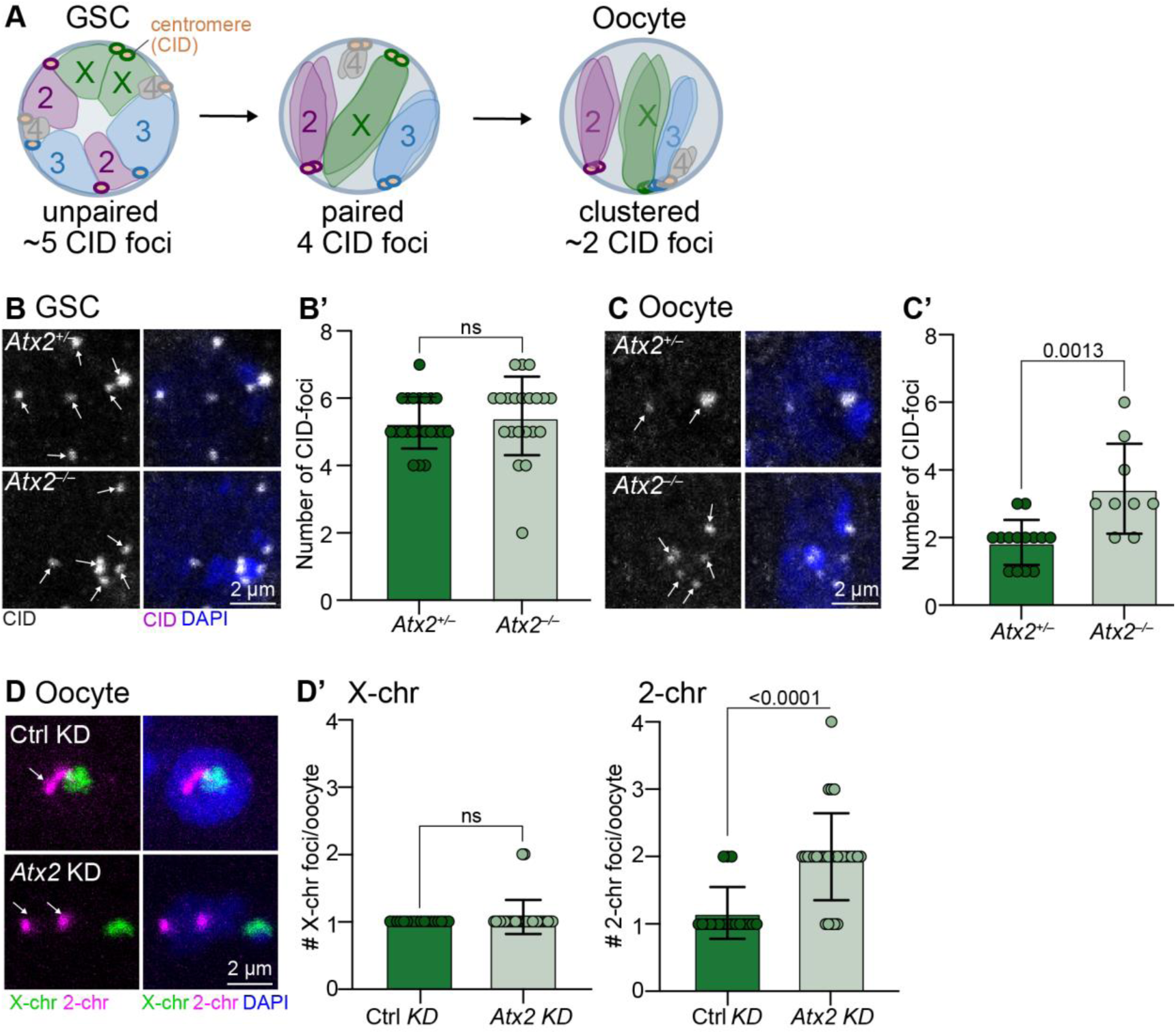
Atx2 is promotes homologous chromosome pairing and clustering. **(A)** Cartoon illustrating chromosome pairing and clustering in germ cells adapted from (Joyce et al. 2013). In GSCs, homologous chromosomes remain unpaired except at the centromeric regions of the X chromosome. By the time the oocyte is specified, all homologs have paired, and clustered apart from chromosome 2. CID marks the centromeric regions of the chromosomes. **(B)** Confocal micrographs of GSCs from heterozygous control and germline ablated *Atx2* ovaries stained with anti-CID to mark centrosomes and DAPI to mark DNA. White arrow indicated CID foci. Scale bar represents 2 µm. **(B’)** Quantification of (B). Data are means ± s.d. of one independent replicate (10 germaria with 2-3 GSCs per replicate). *p*-values, Mann-Whitney non-parametric test. (**C)** Confocal micrographs of pro-oocytes from heterozygous control and germline ablated *Atx2* ovaries stained with anti-CID to mark centrosomes and DAPI to mark DNA. Pro-oocytes were identified by co-staining with anti-Orb. White arrows indicated CID foci. Scale bar represents 2 µm. **(C’)** Quantification of (C). Data are means ± s.d. of one independent replicate (8-10 germaria with 1-2 pro-oocytes per replicate). *p*-values, Mann-Whitney non-parametric test. **(D)** Confocal micrographs of germaria expressing control (mCherry) shRNA or *Atx2* shRNA in the germline stained. FISH was performed with probes against pericentromeric regions of the X- and 2-chromosome. DAPI was used to mark DNA. Oocytes were detected by co-staining with anti-Orb serum (no shown). White arrows indicated 2^nd^ chromosome foci. Scale bar, 2 µm. (**D’**) Quantification of (E). Data are means ± s.d. of one independent replicate (18-30 germaria). *p-values,* Mann-Whitney non-parametric test.

To assess centromere clustering, we used an anti-CID antibody to mark the Drosophila homolog of CENP-A, a histone H3 variant found only at centromeres (Blower and Karpen 2001), and anti-Orb antibody to mark the oocyte (Figure 4A to D). Diploid Drosophila cells have eight chromosomes, which appear as four CID foci when all are paired, and two foci after clustering as occurred (the second chromosome does not cluster with the rest of the chromosomes) (Christophorou et al. 2013; Joyce et al. 2013). As expected from previous work (Christophorou et al. 2013; Joyce et al. 2013), germline stem cells (GSCs) contained an average of five to six CID foci (Figure 4A and B), consistent with the absence of meiotic pairing. By region 2b, however, meiotic germ cells displayed an average of two CID foci, reflecting complete centromere pairing and clustering. In contrast, *Atx2*-depleted germ cells showed a significant increase in CID foci, indicating defective clustering (Figure 4C). In some cases, the number of foci reached as high as six, suggesting that clustering and pairing had completely failed to occur. Together, these results suggest that Atx2 is required for normal centromere pairing and clustering, likely through its role in promoting the expression of SC components.

We next examined the pairing of individual centromeres. Using FISH probes against the pericentromeric regions of chromosome II, which requires SC components for pairing (Christophorou et al. 2013; Joyce et al. 2013), and chromosome X, which pairs independently of the SC and prior to its formation (Christophorou et al. 2013; Joyce et al. 2013), we assessed pairing status (Figure 4D). We again marked oocytes with anti-Orb. We scored the centrosomes as paired when only a single focus was visible or when two foci were separated by less than 0.7 µm, as described previously (Christophorou et al. 2013; Joyce et al. 2013). In control ovaries, nearly all homologous centromeres of the X and II chromosomes were paired, with only a single focus observed on average for both the X and II (Figure 4D). In *Atx2* knockdowns, most chromosome II centromeres failed to pair (Figure 4D), whereas pairing of the X chromosome centromeres remained unaffected (Figure 4D), consistent with the X pairing independently of SC. In *Atx2* knockdowns, chromosome II sister chromatid pairing was also occasionally perturbed, resulting in three or even four distinct foci (Figure 4D). Thus, Atx2 is required for both homologous centromere pairing and clustering, most likely through its positive regulation of the SC.

## DISCUSSION

Here we report a critical role for Atx2 in meiotic progression in *Drosophila*. We show that Atx2 positively regulates key meiotic products, particularly those encoding SC components. Loss or depletion of *Atx2* leads to reduced and impaired SC assembly and maintenance, which in turn causes defects in homologous chromosome pairing and clustering, processes that are essential for normal chromosome segregation during meiosis I and for ensuring the inheritance of a complete complement of chromosomes.

Despite its importance, regulation of the meiotic program remains poorly understood in animals. Although several transcriptional regulators of meiotic entry have been identified in metazoans, such as IME1 in budding yeast and Stra8 in mammals (Kassir et al. 1988), far less is known about how meiotic gene expression is controlled post-transcriptionally. This layer of regulation is critical because both the timing and dosage of meiotic factors must be tightly coordinated. For example, mis-regulation of SC components can lead to the formation of polycomplexes, aberrant SC structures that disrupt gametogenesis (Goldstein 1987; Hughes and Hawley 2020). Our identification of Atx2 as a post-transcriptional inducer of the meiotic program therefore provides an important advance toward understanding how meiotic entry and progression are regulated in animals.

The mechanism by which Atx2 upregulates key meiotic transcripts, including SC components, presents an important avenue for future investigation. Given Atx2’s well described role in post-transcriptional regulation, the simplest model is that Atx2 directly binds these RNAs, stabilizing them and possibly enhancing their translation via its PAM2-mediated interaction with poly(A)-binding proteins (PABPs) and/or by recruiting poly(A) polymerase (Mangus et al. 1998; Satterfield and Pallanck 2006; Yokoshi et al. 2014; Lee et al. 2017; Inagaki et al. 2020; Nadimpalli et al. 2022). However, it remains possible that Atx2 may also or instead act indirectly, for example by modulating other regulators of SC expression. Polycomb group proteins, for instance, have recently been implicated in SC gene transcription, raising the possibility that Atx2 influences SC expression through Polycomb (Feijao T et al. 2022). However, our RNA sequencing analysis did not identify Polycomb components as Atx2 targets. Other proteins, including the SCF ubiquitin ligase complex as well as various kinases and phosphatases, have also been implicated in post-translational regulation of the SC (Williams and Hawley 2025). It remains possible that Atx2 regulates their translation, although we likewise failed to detect these as Atx2 targets in our RNA-sequencing dataset. Thus, further work will be required to define the direct targets through which Atx2 positively regulates the SC, as well as other meiotic proteins such as those involved in recombination.

In sum, our work identifies Atx2 as a central post-transcriptional regulator of meiotic gene expression, particularly SC components. These findings uncover an unexpected layer of meiotic regulation and highlight the importance of RNA-based control mechanisms in safeguarding chromosome segregation and fertility. Clarifying these mechanisms is essential both for fundamental biology and for understanding the origins of human reproductive disorders, as meiotic segregation errors are the primary cause of miscarriage.

## ACKNOWLEDGMENTS

We thank Prashanth Rangan for helpful experimental suggestions and comments on the manuscript. We thank Bloomington Drosophila Stock Center (NIH P40 OD018537) for Drosophila stocks. We obtained the C(3)G and Cid antibodies from R.S. Hawley. The orb 4H8 antibody developed by P. Schedl was obtained from the Developmental Studies Hybridoma Bank, created by the NICHD of the NIH and maintained at The University of Iowa, Department of Biology, Iowa City, IA 52242. This work was supported by the Canadian Institutes of Health Research (FRN 159510) and T.R.H. is part of the University of Toronto Medicine by Design initiative, which receives funding from CFREF. V.L.M. is supported by an Ontario Graduate Scholarship. C.R. is supported by an Irish Research Council Postgraduate Award and B.B. by a DBT-Wellcome Trust India Alliance [IA/I/19/1/504286], and M.R. gratefully acknowledges support from a Research Ireland Future Frontiers Award.

## AUTHOR CONTRIBUTIONS

Conceptualization, V.L.M., C.S., and T.R.H.; methodology, V.L.M., Z.W., J.R.S; reagents, V.L.M., C.R., B.B, and M.R.; investigation, V.L.M, Z.W., C.S and T.R.H.; writing – original draft, V.L.M, and T.R.H ; writing – review & editing, V.L.M, Z.W., J.F.S., C.R., B.B., M.R., C.S., and T.R.H; funding acquisition, V.L.M., and T.R.H.; supervision, T.R.H and C.S..

## DATA AVAILIBILTY STATEMENT

Strains generated in this study are available from the authors upon reasonable request. RNA sequencing data have been deposited in the NCBI GEO database (https://www.ncbi.nlm.nih.gov/geo/info/seq.html), accession number GSE318364. The authors affirm that all other data necessary for confirming the conclusions of the article are present within the article, figures, tables and supplementary materials.

## METHODS

### Drosophila

All fly stocks were reared at 25 °C with controlled humidity on standard medium (cornmeal, agar, yeast and molasses). All stocks used are listed in Table S3. Genotypes for each figure are listed in Table S4. All knockdown and mutant experiments were conducted at 25 °C. We used FlyBase (release FB2025_05) (Öztürk-Çolak et al. 2024) to obtain stock, expression and sequence information.

### Generation of Drosophila strains

N-terminally GFP-tagged Atx2 and FRT-Atx2::GFP-FRT flies were generated using CRISPR genome engineering and homologous recombination (HR), as previously described (Port et al. 2014; Bakthavachalu et al. 2018). In summary, a pair of gRNAs were designed to target the 5’UTR and 3’UTR of Atx2 from a single plasmid (pU6.2 plasmid). To create the N-terminally GFP-tagged Atx2 donor plasmid, the whole *Atx2* gene was cloned into the Puc19 background, and the GFP tag was inserted just before the start codon to. To create the FRT-Atx2::GFP-FRT donor plasmid, GFP was cloned to the C-terminus of the *Atx2* gene, and FRT sequences were inserted on both sides of the coding sequences into the PUC19 plasmid. To facilitate the screening of the recombinant animals, a *dsRed* sequence driven by the *3XP3* promoter was inserted into the first intron of the *Atx2* gene, transcribed in the opposite direction of the *Atx2* gene. The gRNA and HR donor plasmids were injected into a *Act5c-Cas9-Lig4* strain (N-terminally GFP tagged Atx2) and *nos-Cas9* strain (FRT-Atx2::GFP-FRT) by BestGene Inc., and transgenics were screened for DsRed expression in the eyes. The established strains were then validated by sequencing.

### Fluorescence activated cell sorting (FACS)

The ovary dissociation protocol was adapted from Vallés and Huynh (2020) and Slaidina et al. (2021). Briefly, 170 – 200 control knockdown and 200 – 300 *Atx2* knockdown young ovaries (< 24 hours old) were dissected in Schneider’s Drosophila media supplemented with 10% FBS. Ovaries were dissociated in cell dissociation buffer (ThermoFisher Scientific, 13150016) with collagenase (0.25 mg/mL; Gibco, 17018029) and elastase (0.4 mg/mL; BioShop Canada, ELA292) at 37°C for 30 min with shaking. Dissociated cells were run through 70 µm (Genesee Scientific, 25-376) and 30 µm (Miltenyi Biotec, 130-041-407) cell strainers and resuspended in PBS with 1% BSA prior to sorting. Propidium iodide (3 µM; BioShop Canada, PPI777) was used to measure identify and sort for live cells. The BD FACSymphony S6 SE cell sorter was used to isolated GFP positive cells. Gating conditions to exclude GFP-negative cells were set using dissociated *w1118* ovary cells as negative controls.

### RNA extraction and RT-qPCR

Total RNA was extracted from sorted cells according to manufacturer’s protocol using TRI-Reagent, chloroform (Sigma-Aldrich, 472476), GlycoBlue (ThermoFisher Scientific, AM9516), 2-propanol (Sigma-Aldrich, I9516) and ethanol, and eluted in nuclease-free water. Purified RNA was treated with Turbo DNAse (ThermoFisher Scientific, 2238G2). An additional ethanol precipitation was carried out according to Green and Sambrook (Green and Sambrook 2020).

For RT-qPCR, 40 ng of total RNA was used with SuperScript IV Reverse Transcriptase (ThermoFisher Scientific, 18090050) and <u>random hexamers (ThermoFisher Scientific, SO142)</u> to make cDNA. Quantitative PCRs for stage specific markers were carried out on 1/4 of reverse transcription reaction and 300 nM of each primer pair using the SensiFAST SYBR No-ROX kit (FroggaBio, BIO-98050) and a Bio-Rad CFX384/C1000 Touch system (Bio-Rad). The PCR program was as follows: 2 min at 95 °C; 45 cycles of 95 °C for 5 s and 60 °C for 30 s. Results of qPCRs for transposable elements were normalized to the mean of value obtained of CG2698 and Und. Results were calculated using the following formula: ΔΔCt = 2-(ΔCt*^Atx2^* ^RNAi^ – ΔCt*^mCherry^* ^RNAi^), where ΔCt = Ct (gene) – Ct (mean of CG2698 and Und). Data was plotted using GraphPad Prism 10.

Primers used were as follows:

*CG2698* (CTTCAGCATTTGTGGCAGAC; ATGTGTCGCTCTGGTGACTG),

*Und* (GCAAGAAAAGCGGTCAGACT; CGTGTTGATACGGTCCAGAG),

*bam* (GGGAGGTCCGATCTATTGCG; CGATCAGAGCGGAGAGGAAC),

*Orc1* (ACGACTTTCATGGGCGTCTA; TCTGAGGGCGTAGTGAATGT),

*CG15628* (TGGCCCGACAAACCGATAAT; GAGTCGTTCCATCACACCGA),

*Nlg2* (GCAGTCCTCGGATCAGATGT; ATGTACACGTTTCTTGCCGC),

*mnd* (ATCTCAATGCCGGTGGTCAC; GTGACCATTCCTTGCACCGA), and

*slif* (CTGAACACATGGATCCGCTT, TCGGAGAACTCGTAGGGTC).

### RNA sequencing and analysis

Purified RNA concentration was measured by Qubit 4 Fluorometer and quality was measured by Agilent 2100 Bioanalyzer by The Centre for Applied Genomics (TCAG) at The Hospital for Sick Children. Ribosomal RNA (rRNA)-depleted cDNA library preparation was performed by TCAG using QIAseq FastSelect –rRNA Fly Kit (Qiagen, 333262) and NEBNext Ultra II Directional RNA Library Prep Kit (NEB, E7760) without poly(A) selection with 18 ng of RNA. TCAG performed sequencing using a S4 flowcell on an Illumina NovaSeq 6000 to generate 150-bp paired-end reads, and at an average depth of 100 million reads.

Libraries were evaluated for quality using FastQC v0.11.9 (RRID:SCR_014583) (Andrews, 2019). Adaptor sequences were trimmed using default settings of Trim Galore v0.6.7 (RRID:SCR_011847) (Krueger, 2021) which is a wrapper around Cutadapt v2.10 (RRID:SCR_011841) (Martin, M., 2011). Residual rRNA sequences were removed using RiboDetector v0.2.7 using default settings for paired-end reads (Deng, Z. et al., 2022) or with SortMeRNA 4.3.6 and rRNA sequences documented by SILVA (Kopylova et al. 2012; Quast et al. 2013). Transcript level read counts were obtained using Salmon v1.9.0 (RRID:SCR_017036) (Patro et al., 2017) by indexing to dmel r6.54 chromosome and transcripts fasta sequences hosted on FlyBase (Gramates et al., 2022). Genes with less than 1 TPM in total across control samples were filtered out as inactive genes. Fifty-three contaminating chorion and vitelline membrane genes (GO term IDs: 0032529, 0060388, 0042600, 0007305) were excluded at the filtering stage. Sample normalization using the median-ratios normalization method and differential expression analysis was carried out using the DESeq2 R package v1.42.0 (RRID:SCR_015687) (Love et al., 2014). Genes were considered differentially expressed if the Benjamini-Hochberg adjusted p-value was less than 0.05 and the absolute log_2_ fold change was greater than or equal to 1.5. Volcano plots were generated by ggplot2 v3.4.4 (RRID:SCR_014601) (Wickham, 2016). GO-term enrichment analysis was performed using gProfiler2 v0.2.2 (RRID:SCR_018190) (Kolberg et al., 2020) using the respective background, false-discovery rate correction method and excluding GO electronic annotations. R v4.3.1 was used to perform the analysis. Box plots were generated using ggplot2 v3.5.2 (RRID:SCR_014601) (Wickham, 2016), ggbreak v0.1.5 (RRID: SCR_024824) (https://github.com/YuLab-SMU/supplemental-ggbreak), and ggpubr v0.6.1 (RRID: SCR_021139) (https://rpkgs.datanovia.com/ggpubr/).

### Ovary immunofluorescence

Adult ovaries were stained according to standard procedures. Briefly, ovaries from yeasted flies were dissected in PBS and fixed in 4% formaldehyde (Thermo Scientific, 28908) in PBS for 15 min. Ovaries were then permeabilized with 1% Triton X-100 (BioShop Canada, TRX506) in PBS for 20 min. Ovaries were incubated with primary antibodies diluted in 1% PBST (1% (w/v) bovine serum albumin [BSA, BioShop Canada, ALB001], 0.1% Triton X-100, PBS) overnight at 4°C followed by incubation with the appropriate secondary antibodies diluted in 1% PBST for 2 hours at room temperature. Ovaries were counterstained with 4′,6-diamidino-2-phenylindole (0.2 – 1 µg/mL; Cell Signaling Technologies, 4083S) for 15 min and/or phalloidin Alexa Fluor 633 (1:250; Invitrogen, A22284) for 2 hours with secondary antibodies. Ovaries were mounted in VECTASHIELD Antifade Mounting Medium (BioLynx, VECTH1000). All images were acquired with a Leica SP8 inverted scanning confocal microscope using 63x (NA 1.4, immersion oil) objectives. All experiments were performed using multiple sections (z-stacks) from confocal images. Image analysis and maximum projections was performed using FIJI (Schindelin et al., 2012). Confocal images shown are maximum intensity projections of three optical slices (0.3 µm each), except for Figure 1E (21 slices, 0.3 µm each) to capture the full width of the germarium, and Figure 4B (14 slices, 0.3 µm each) and 4C (8 slices, 0.3 µm each) to capture the full nuclei.

The following primary antibodies were used: mouse anti-orb 4H8 (1:100; deposited to the DSHB by P. Schedl); mouse anti-C(3)G and rat anti-CID (1:500 each; gifts from R.S. Hawley); and rabbit anti-GFP (1:1000; Invitrogen, A11122). The following secondary antibodies were used at a 1:500 dilution: donkey anti-mouse Cy3 (Jackson ImmunoResearch Labs, 715-165-151); donkey anti-rabbit Alexa Fluor 647 (Jackson ImmunoResearch Labs, 711-605-152); goat anti-rabbit Oregon Green 488 (Invitrogen, O-11038); goat anti-mouse Alexa Fluor 488 (Invitrogen, A-11001); donkey anti-rat Cy3 (Jackson ImmunoResearch Labs, 712-165-153).

### Hybridization Chain Reaction (HCR) RNA Fluorescence In situ Hybridization

Probes were ordered from Molecular Instruments, Inc and protocol was adapted Slaidina et al. (2021) and Molecular Instruments’ sample-in-solution protocol. Adult ovaries from yeasted flies were dissected in PBS and fixed with 4% formaldehyde and 0.1% Tween-20 (Sigma-Aldrich, P9416) in PBS for 20 min at room temperature. Ovaries were washed twice in 0.1% Tween-20 in PBS post-fixation for 5 min and dehydrated with sequential washes of 25%, 50%, 75% and 100% ethanol (Commercial Alcohols, 22734) in PBS on ice. Ovaries were stored at -20°C at least overnight and up to 7 days, and were rehydrated with sequential washes of 100%, 75%, 50% and 25% ethanol in PBS on ice. Ovaries were permeabilized for 2 hours with 1% Triton X-100 in PBS at room temperature and fixed with 4% formaldehyde and 0.1% Tween-20 in PBS for 20 min. After fixation, ovaries were washed twice on ice with 0.1% Tween-20 in PBS, twice with 0.1% Tween-20 in 0.5x PBS and 2.5x SSC buffer, and then twice 0.1% Tween-20 in 5x SSC buffer. Samples were incubated on ice with 200 µL probe hybridization buffer for 5 min followed by pre-hybridization for 30 min at 37°C. Ovaries were hybridized with the 8 pmol of probes overnight in 200 µL pre-warmed hybridization buffer at 37°C. After hybridization, ovaries were washed four times with pre-warmed probe wash buffer for 15 min each at 37°C and twice with 0.1% Tween-20 in 5x SSC, and then equilibrated in 500 µL of amplification buffer at room temperature for 5 min. Meanwhile, Alexa Fluor 594 hairpin pairs were prepared by heating 30 pmol of each hairpin separately 95°C for 90 seconds and cooling to room temperature in the dark for 30 min. The hairpin pairs were subsequently added to ovaries in amplification buffer and incubated in the dark overnight at room temperature. Following probe amplification, ovaries were washed with 0.1% Tween-20 in 5x SSC twice for 5 min, twice for 30 min and once for 5 min. Primary and secondary antibody staining was carried out as described above in the dark.

### C(3)G protein and mRNA Quantification

C(3)G protein and mRNA were quantified using FIJI. For mRNA, the region of interest (ROI) was drawn around the germarium of a sum intensity projection of 41 slices (0.3 µm each) encompassing the germarium. For protein, the ROI was drawn around the single oocyte nuclei at region 2B with a sum intensity projection of 10 slices (0.3 µm each). Fluorescence intensity was calculated using the corrected total cell fluorescence method (Gavet and Pines 2010) where the mean background used was the average of mean fluorescence intensity of two somatic cells encapsulating an egg chamber. Each value was normalized to the mean of control values to obtain mRNA/protein fold change. Data was plotted using GraphPad Prism 10.

### DNA FISH

Protocol was adapted from McKim et al. (2009) (McKim et al. 2009). Briefly, 4% formaldehyde-fixed ovaries were rinsed with 2X SSCT (0.1% Tween-20) twice. Ovaries were washed sequentially for 15 min with 2X SSCT + 20% formamide (VWR 97062-006), 40% formamide and 50% formamide at room temperature and nutation. After final wash, ovaries were incubated in fresh 2X SCCT/50% formamide at 37°C for 1-2 hours. After removal of 2X SSCT/50% formamide, ovaries were incubated in 40 µL of hybridization solution (10% dextran sulfate [BioShop DEX001.50], 50% formamide, 2X SSC, 1 mM EDTA and 1 µM of each probe, sequences below) at 91°C for two min, mixed and then subsequently incubated at 37°C overnight in the dark. Pre-warmed (37°C) 2X SSCT/50% formamide was added to dilute dextran sulfate and allow ovaries to settle before washing three times with pre-warmed 2x SSCT/50% formamide for 20 min each in the dark. The formamide concentration was brought down by gradually washing with 2X SSCT/40% formamide and 20% formamide for 10 min each. Ovaries were washed twice with 2X SSCT and finally with PBSTB (0.1% Triton X-100 and 1% BSA) before blocking with PBSTB for 1 hour at room temperature before continuing with immunofluorescence using anti-Orb, as described above. Images were acquired with a Leica SP8 inverted scanning confocal microscope using 63x (NA 1.4, immersion oil) objective. Confocal images shown are maximum intensity projections of 26 optical slices (0.3 µm each).

X-chromosome rRNA probe (Lu Kevin L 2018):

Cy3 5’ CCACATTTTGCAAATTTTGATGACCCCCCTCCTTACAAAAAATGCG

Chromosome 2 repeat probe (Bhargava et al. 2020):

Cy5 5’ AACACAACACAACACAACACAACACAACAC

## FIGURES

**Supplementary Figure 1.**
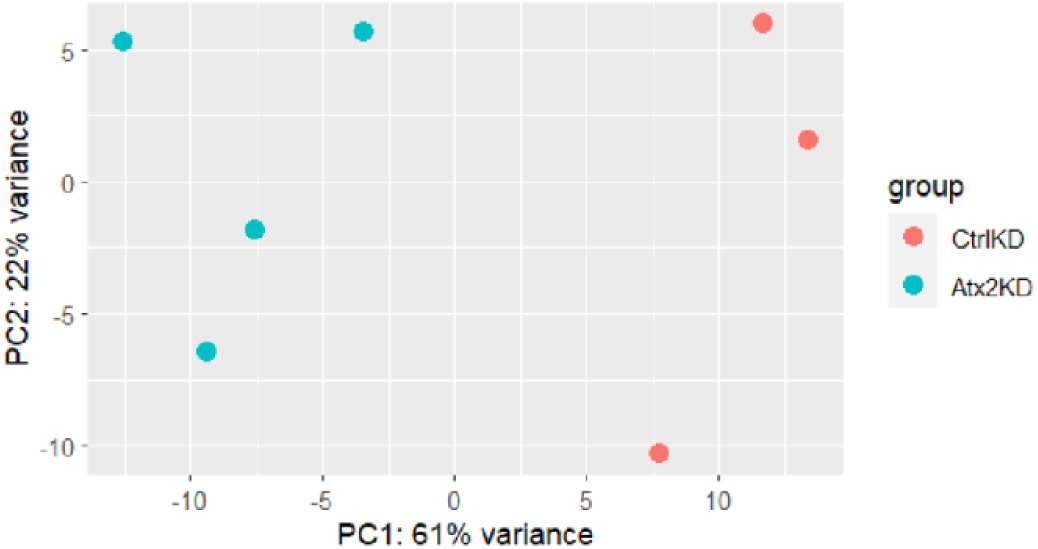
Principal component analysis of control and *Atx2* knockdown RNA sequencing experiments.

**Supplementary Table 1.** Differential transcript expression in *Atx2* knockdown vs Control knockdown meiotic germ cells. Normalized fold change presented between control vs Atx2 KD meiotic germ cells.

**Supplementary Table 2.** GO term analysis of transcript downregulated in Atx2 knockdown meiotic germ cells.

**Supplementary Table 3.** Drosophila strains used.

**Supplementary Table 4.** Drosophila genotypes for each figure.

